# Preference-Independent Encoding of Visual Saliency Within but Not Cross Features in the Mouse Superior Colliculus

**DOI:** 10.1101/2024.03.04.583246

**Authors:** Ruixiang Wu, Jinhuai Xu, Chunpeng Li, Zhaoji Zhang, Ling-yun Li, Ya-tang Li

**Affiliations:** Chinese Institute for Brain Research, Beijing 102206, China; College of Biological Sciences, China Agricultural University, Beijing 100193 China; College of Chemistry and Molecular Engineering, Peking University, Beijing 100091, China; Department of Neurobiology, School of Basic Medical Sciences, Capital Medical University, Beijing 100069, China

**Keywords:** superior colliculus, visual saliency, two-photon calcium imaging, preference-independent encoding

## Abstract

Detecting conspicuous stimuli in a visual scene is crucial for animal survival, yet it remains debated how the brain encodes visual saliency. Here we investigate how visual saliency is represented in the superficial superior colliculus (sSC) of awake mice using two-photon calcium imaging. We report on a preference-independent saliency map in the sSC. Specifically, salient stimuli evoke stronger responses in both excitatory and inhibitory neurons compared to uniform stimuli, with similar encoding patterns observed in both neuron types. The largest response occurs when a salient stimulus is positioned at the receptive field center, with contextual effects extending ∼40° away from the center. The response amplitude correlates well with the saliency strength of stimuli and is not influenced by the orientation or motion direction preferences of neurons. However, saliency encoding does depend on specific visual features. Furthermore, neurons involved in saliency encoding exhibit weak orientation or direction selectivity, suggesting a complementary relationship between the saliency map and the feature map.

## 1 Introduction

Visual attention plays an essential role in sifting through massive visual information that enters the eyes, a critical factor for animals to survive in a complex and ever-changing environment [1]. Visual attention has two types: endogenous (goal-driven/top-down) attention and exogenous (scene-driven/bottom-up) attention. Exogenous attention is naturally directed towards salient objects that “pop out” in the scene [2, 3, 4, 5, 6].

Saliency, defined as the conspicuousness of a specific visual area within a scene, can arise from various physical attributes such as orientation, color, motion direction, size, and shape. It is still under debate where and how visual saliency is encoded in the brain. A theoretical model proposed that the brain has a saliency map [7, 8]. This map integrates inputs from diverse feature maps and represents saliency strength independent of specific features. This saliency map merges with goal-driven attention to generate a priority map, ultimately guiding where to attend [9, 10]. Multiple brain regions in primates have been identified as encoding visual saliency, including the primary visual cortex (V1) [11, 12, 13, 14], the superficial and intermediate layers of the superior colliculus (sSC and iSC) [15], V4 [16], posterior parietal cortex [17], and the frontal eye field [18]. However, with the exception of the sSC and V1, these regions also process goal-driven attention, suggesting that they might inherit the saliency map from upstream regions and host a priority map [19].

While the model suggests that ideal saliency-encoding neurons respond to salient stimuli independently of their preference within a specific visual feature or cross different features, there is a lack of conclusive experimental evidence from both V1 and sSC. In the V1 of both anesthetized and awake cats and monkeys, salient stimuli made by orientation have elicited larger, similar, or smaller responses compared to iso-orientation stimuli [12, 13, 20, 21, 22]. Similar inconsistencies have also been observed in neuronal responses to salient stimuli made by motion direction [14, 23, 24]. However, these contradictory findings can be largely reconciled by considering a neuron’s feature preference. That is, a neuron might be more responsive to certain orientations or motion directions. This suggests that V1 neurons respond to salient stimuli made by the same feature in a preference-dependent manner [12]. The saliency encoding in the sSC is also in debate. While recent work in awake monkeys suggests a correlation between neural activity and saliency strength, the role of feature preference remains unsolved [15]. Additionally, studies in mice complicate the picture, as saliency encoding in the sSC is affected by anesthesia [25, 26]. Other factors like cortical integrity and the number of neurons recorded could also influence data interpretations.

In the present work, we carried out *in vivo* two-photon calcium imaging in the sSC of head-fixed awake mice with an intact cortex. We aim to understand how the encoding of visual saliency is affected by different visual features and contributed by excitatory and inhibitory neurons at different depths of the sSC. To achieve this, we expressed non-floxed GCaMP8 in the mouse sSC where GABAergic or glutamatergic neurons were genetically labeled by tdTomato. We then measured and analyzed neuronal responses to salient stimuli based on either orientation or motion direction. Our results demonstrate that throughout the mouse sSC, excitatory and inhibitory neurons encode visual saliency in a similar way. This encoding is independent of their preference for specific orientation or motion direction, but it does depend on the particular feature that creates the saliency.

## 2 Results

### 2.1 Excitatory and inhibitory neurons respond strongly to salient stimuli

To investigate how visual saliency is encoded in mouse sSC, we applied two-photon calcium imaging in head-fixed awake mice (Fig. 1A and B) [27]. We measured the activity of both excitatory and inhibitory neurons in response to visually salient stimuli (Fig. 1C). We imaged the posterior-medial SC while leaving the cortex intact as previously described [28, 29]. Across 54 image planes and up to a depth of 300 *µ*m, we recorded a total of 2565 excitatory neurons and 4455 inhibitory neurons in seven animals. Throughout the imaging session, the animal exhibited occasional locomotion and eye movements (Fig. S1A and D). During periods of locomotion, we observed a positive correlation between pupil size and locomotion speed, and neuronal responses showed either a negative or positive correlation with pupil size (Fig. S1B and C), consistent with previous findings [30].

**Figure 1.**
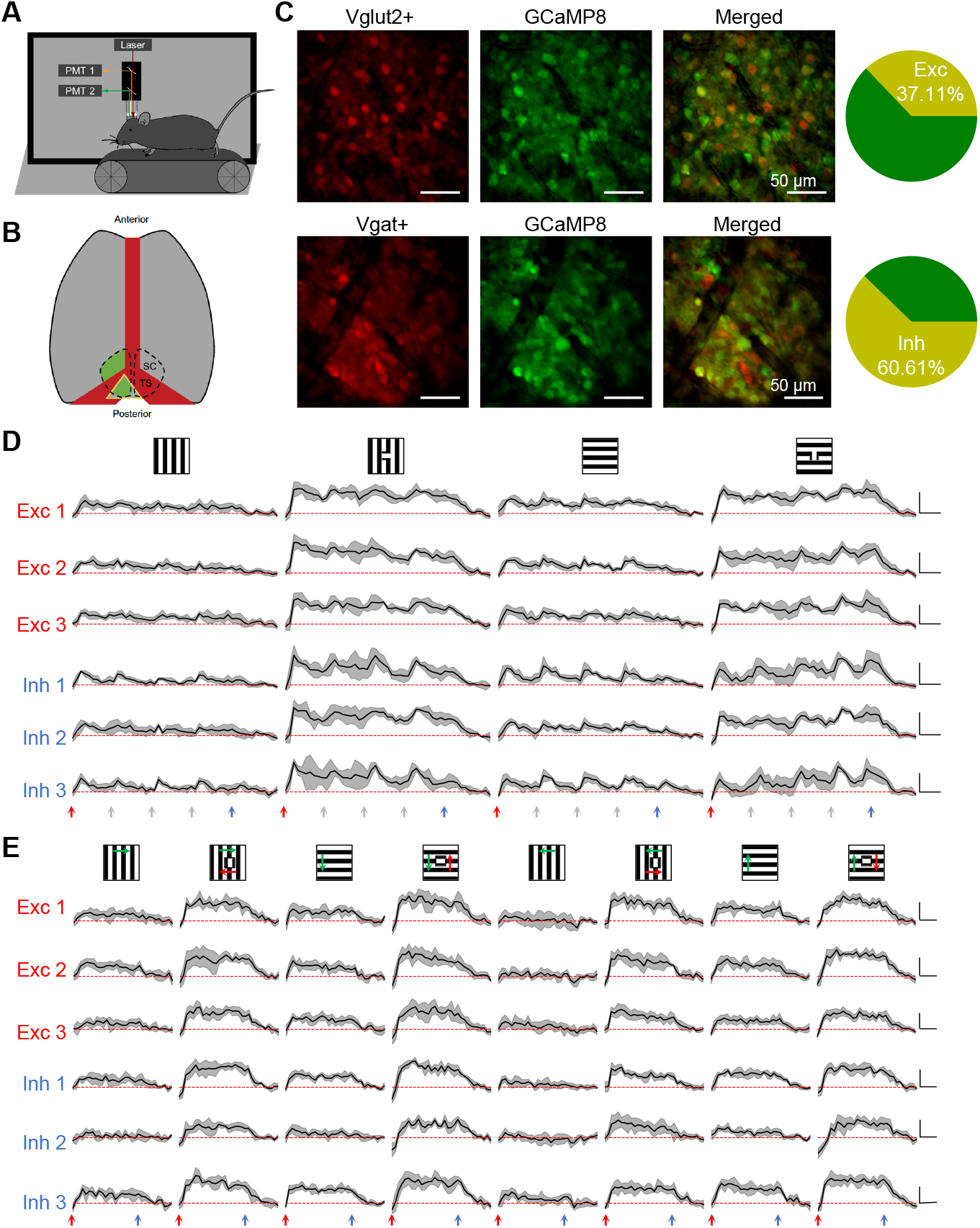
Two-photon calcium imaging reveals robust visual responses to salient stimuli in the awake mouse superior colliculus. **A**. Schematic of the experimental setup. Mice were head-fixed and free to run on a treadmill. Visual stimuli were presented on a screen. Neuronal calcium activity was imaged using two-photon microscopy. PMT, photomultiplier tube. **B**. Schematic of mouse brain anatomy after insertion of a triangular transparent plug to expose the posterior-middle portion of the superior colliculus beneath the posterior cortex. TS: transverse sinus. **C**. A mean projection of tdTomato-labeled Vglut2+ or Vgat+ neurons and GCaMP8+ neurons in two mice. The right panel shows the proportion of double-labeled neurons in both Vglut2-tdTomato and Vgat-tdTomato mice. **D**. Example response profiles of excitatory and inhibitory neurons to salient flashing gratings and backgrounds. Gray shade indicates the SD across 10 trials. Blue and red arrows mark the onset and offset of visual stimuli, and gray arrows mark the phase shift of the gratings. Scale: 30% ΔF/F_0_, 0.5 s. **E**. Example response profiles to salient moving gratings and backgrounds. Gray shade indicates the SD across 10 trials. Blue and red arrows mark the onset and offset of visual stimuli. Scale: 30% ΔF/F_0_, 0.5 s.

Excitatory and inhibitory neurons were distinguished by expressing tdTomato in either Vglut2+ or Vgat+ neurons (Fig. 1C). In both lines, the percentage of excitatory neurons was 40%, in line with the fluorescence *in situ* hybridization data [31]. To examine how neurons encode visual saliency, we measured their calcium responses to salient flashed gratings (SFG, Fig. 1D) or salient moving gratings (SMG, Fig. 1E) at the RF center with different backgrounds. Both excitatory and inhibitory neurons show stronger responses to salient stimuli compared to the background alone, suggesting an important role of the sSC in encoding visual saliency.

### 2.2 Neural encoding of visual saliency is independent of orientation preference

Neurons in sSC are retinotopically organized and show orientation preference. A neuron’s response to SFGs is determined by three factors: 1) the distance between its RF center and the SFG; 2) the orientation contrast between the target and the background; and 3) its orientation preference. To understand how these three factors affect neuronal responses, we measured RFs and orientation preferences of individual neurons, along with their responses to SFGs at different spatial locations relative to RFs. For both excitatory and inhibitory neurons, the strongest response is elicited by the SFG at the RF center, which gradually decreases as the SFG moves away (Fig. 2A, 2B, S2A). This observation suggests that these collicular neurons function as saliency detectors.

**Figure 2.**
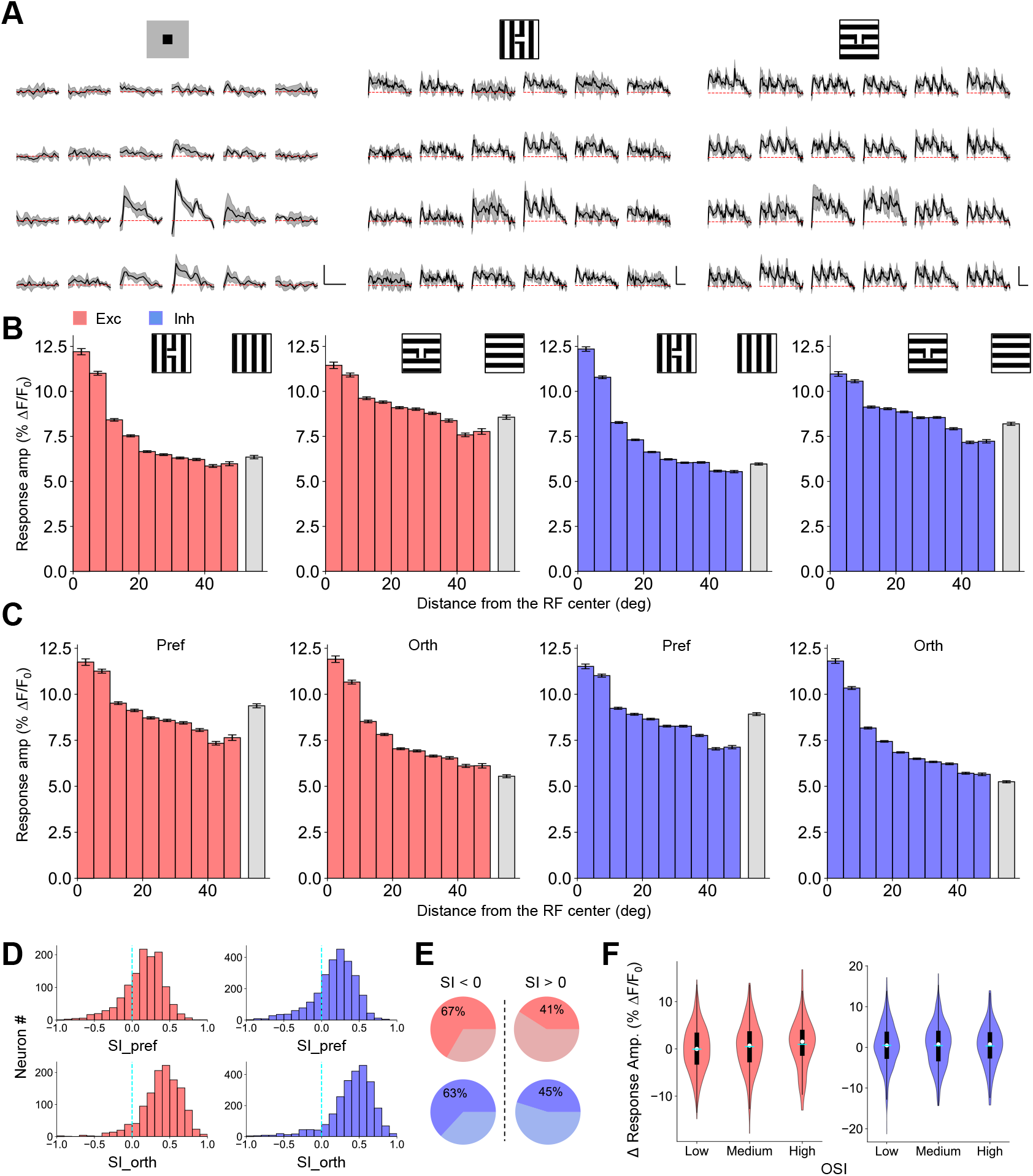
Neural encoding of visual saliency is independent of orientation preference. **A**. Calcium responses of an example neuron to flashed white or black squares of 10° against a grey background, flashed horizontal and vertical gratings of 10° against an orthogonal background. The targets were displayed at each of the 4 × 6 locations. Gray shade indicates the SD across 10 trials. Scale: 30% Δ F/F_0_, 1 s. **B**. Response amplitude to SFGs with the target at different distances from the RF center (colored bars), as well as the amplitude to the background (gray bars) for both excitatory (N=1273) and inhibitory (N=2684) neurons. **C**. Response amplitude to SFGs and backgrounds sorted by orientation preference. **D**. Histograms of SI for excitatory and inhibitory neurons. Cyan dashed lines mark the 0. **E**. The percentage of orientation-selective neurons in both non-saliency-encoding (left) and saliency-encoding (right) populations. **F**. The difference in response amplitude elicited by the preferred and orthogonal orientations for saliency-encoding neurons with different levels of orientation selectivity. Low: OSI*<*0.25; Medium: 0.25 ≤OSI*<*0.5; High: OSI ≥ 0.5. The means of the three groups are the same (one-way ANOVA, *p*=0.11 for excitatory neurons, *p*=0.92 for inhibitory neurons).

To disentangle the contribution of orientation contrast and neurons’ orientation preferences, we sorted SFG-elicited neuronal responses based on whether the neurons prefer vertical or horizontal gratings. For both types of neurons, the response amplitude at the RF center remains constant regardless of whether the background orientation is preferred (Fig. 2C), indicating that saliency encoding in the sSC is independent of the orientation preference. As the target moves further from the RF center, the response amplitude gradually decreases, up to ∼40°. For the preferred orientation, the response amplitude becomes comparable to that evoked by the background when the distance reaches ∼10°, the typical RF size of sSC neurons [32, 33]. A further decline of the amplitude with the distance up to ∼40° indicates a long-range suppression effect from the target. Conversely, a long-range facilitation effect is observed if the background grating is orthogonal to neurons’ preferred orientations. This contrasting long-range effect for preferred and orthogonal orientations could be implemented by local neurons with extended dendritic arbors. Examples include inhibitory horizontal cells and excitatory wide-field cells, which have dendrites that extend up to ∼500 *µ*m [34].

To further confirm the preference-independent saliency encoding, we calculated a saliency index (SI) to quantify each neuron’s contribution to saliency encoding (see Methods). Compared to the background, about 80% neurons show a stronger response to the salient stimuli for both types of neurons, even when the background matched their preferred orientation (Fig. 2D). Interestingly, a smaller proportion of saliency-encoding neurons exhibit orientation selectivity (OS) compared to non-saliency-encoding neurons (Fig. 2E). These neurons show similar response amplitude to salient stimuli regardless of whether the background aligns with or is orthogonal to their preferred orientation (Fig. 2F), supporting the notion that saliency encoding in the sSC is independent of orientation preference. In addition, the saliency encoding remains consistent across different depths within the sSC (Fig. S2B and C).

### 2.3 Neural encoding of visual saliency is independent of direction preference

Saliency in visual stimuli can be induced by different features. To explore whether the orientation preference-independent encoding of saliency extends to other features, we measured and analyzed neuronal responses to SMGs. Similar to SFG-evoked responses, neurons exhibit the strongest responses when the target is at the RF center, and the response amplitude decreases when the target moves away from the RF center (Fig. 3A and B). Further analysis reveals a negative correlation between the encoding of visual saliency and motion direction (Fig. 3C and 3D), as observed for orientation. For neurons that encode both saliency and motion direction, their response amplitude remained similar regardless of whether the background moves in the preferred or null direction (Fig. 3E). These findings suggest that encoding of visual saliency is independent of a neuron’s preference for both orientation and motion direction.

**Figure 3.**
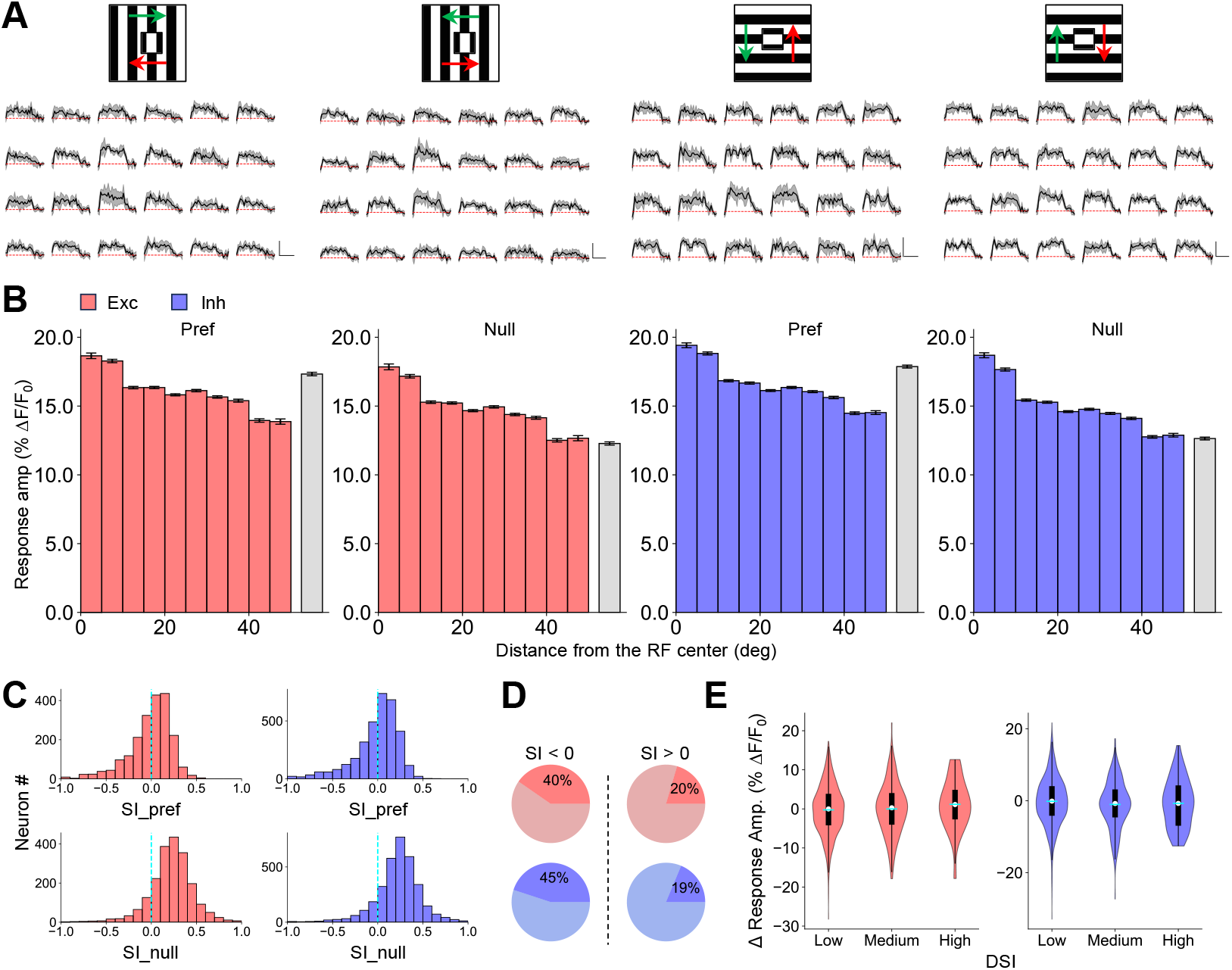
Neural encoding of visual saliency is independent of direction preference. **A**. Calcium responses of an example neuron to moving gratings against a background moving in four directions. Green arrows indicate the motion direction of the background, and red arrows indicate the direction of the target. Gray shade indicates the SD across 10 trials. Scale: 30% Δ F/F_0_, 1 s. **B**. Response amplitude to SMGs (colored bars) and the background (gray bars) for both excitatory (N=2013) and inhibitory (N=3434) neurons, sorted by direction preference. **C**. Histograms of SI calculated from SMG-elicited responses for excitatory and inhibitory neurons. Cyan dashed lines mark the 0. **D**. The percentage of direction-selective neurons in both non-saliency-encoding (left) and saliency-encoding (right) populations. **E**. The difference in response amplitude elicited by the preferred and null directions for saliency-encoding neurons with different levels of direction selectivity. Low: DSI*<*0.25; Medium: 0.25 ≤ DSI*<*0.5; High: DSI ≥ 0.5. The means of the three groups are the same (one-way ANOVA, *p*=0.46 for excitatory neurons, *p*=0.11 for inhibitory neurons).

### 2.4 Neurons in the sSC encode saliency strength

An ideal saliency-encoding neuron wouldn’t just be independent of orientation and direction preference, it would only respond to the saliency strength of a visual stimulus. To examine whether neurons in the sSC meet this criterion, we systematically changed the orientation of the target against a fixed vertical or horizontal background, with an orientation contrast of 90° indicating the maximal saliency strength. Our findings indicate that neuronal responses indeed reflect the orientation contrast rather than the absolute orientation of the target (Fig. 4A and E). To quantify the contributions of orientation contrast, we examined the relationship between neuronal responses and orientation contrast for both excitatory and inhibitory neurons. For saliency-encoding neurons that are not selective to orientation, the response amplitude gradually decreases when the orientation contrast deviates from 90°, indicating the neuronal responses encode the orientation contrast (Fig. 4B and F). The same relationship is observed for OS neurons (Fig. 4C and G). Note that the response amplitude evoked by the SFG with a 30° contrast is either smaller (for excitatory neurons) or comparable (for inhibitory neurons) compared to the response evoked by the preferred background. This finding aligns with the notion that the saliency encoding is independent of orientation and direction preference (Fig. 2 and 3) and corresponds to the findings observed in the V1 of cats and monkeys [14]. This independence is reinforced by averaged SFG-evoked responses aligned with the preferred orientation of each neuron (Fig. 4D and H). Furthermore, the difference in response amplitude between 30° and 90° orientation contrasts only reflects the difference in their saliency strength, and is independent of the neuron’s orientation selectivity (Fig. 4E and I).

**Figure 4.**
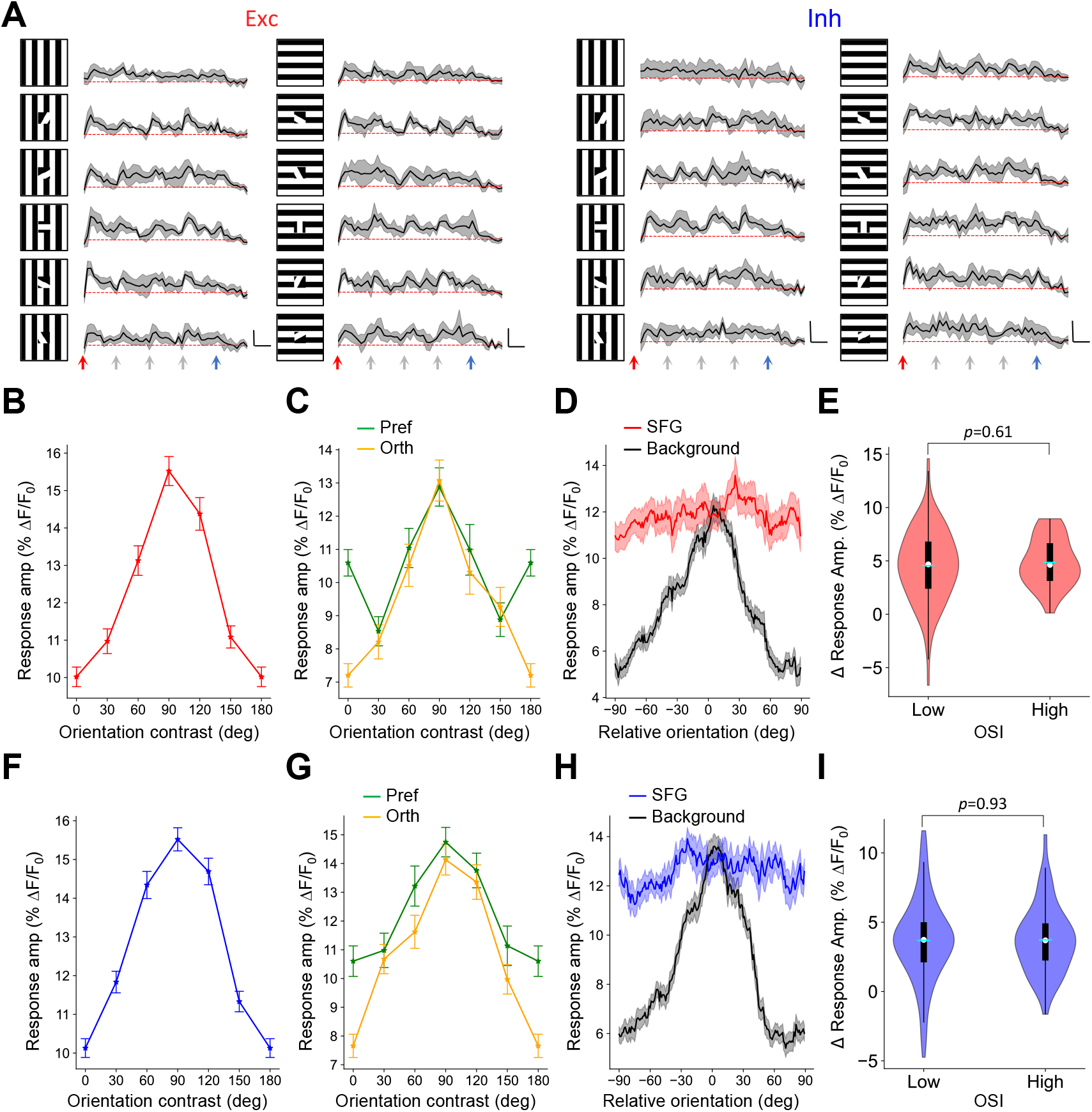
Neurons in the sSC encode saliency strength. **A**. Response profiles of example excitatory and inhibitory neurons to SFGs with 6 different orientation contrasts in two backgrounds. Gray shade indicates the SD across 10 trials. Scale: 30% Δ F/F_0_, 0.5 s. **B**. Averaged calcium responses to different orientation contrasts for neurons with SI>0 and OSI<0.25. Error bars represent SEM, N=98. **C**. Averaged calcium responses in preferred and orthogonal backgrounds for neurons with SI>0 and OSI>0.25, N=36. **D**. The averaged responses evoked by the SFG and the background aligned to the preferred orientation of each neuron, N=134. Shades indicate SEM. **E**. The difference in response amplitude elicited by 30° and 90° contrasts for saliency-encoding neurons with different levels of orientation selectivity. Low: OSI*<*0.25; High: OSI ≥ 0.25, *t*-test. **F-I**. The same plots as **B-E** for inhibitory neurons. For neurons with OSI<0.25, N=116; for neurons with OSI>0.25, N=51.

### 2.5 Encoding of visual saliency depends on specific visual features

We have shown that the response amplitude of saliency-encoding neurons correlates well with the saliency strength of the stimuli and is independent of their preference for orientation or motion direction. Another prediction of the theoretic model is that the response amplitude of neurons should also be independent of their preference for specific visual features that create that saliency (Fig. 5A). Note that while the saliency strength of SFG can be quantified as the orientation contrast, it is difficult to compare the saliency strength between SFG and SMG. Ideally, the difference in response amplitude (ΔR) elicited by two salient stimuli would only depend on the difference in their salient strength, not on the neurons’ preference for specific orientations, motion directions, or features (Fig. 5B). For example, ΔR to two stimuli with the same saliency strength should be close to zero and independent of OSI or DSI (Fig. 2F and 3E, the orange line in Fig. 5B). When the saliency strength is different, ΔR should be constant; this applies to salient stimuli made by the same feature (Fig. 4E and I, the green line in Fig. 5B) and different features (the blue line in Fig. 5B). To test whether saliency encoding is independent of preference for specific visual features, we compared neuronal responses evoked by SFG and SMG. The preference of neurons for specific features was quantified with a feature selectivity index (FSI, see Methods). Our data show a strong positive correlation between FSI and ΔR (Fig. 5C). Dividing the neurons into three groups with different levels of feature selectivity, ΔR is significantly different among these groups (Fig. 5D). Thus, our results demonstrate that the neural encoding of visual saliency depends on the preference for specific visual features, which is against the theoretic prediction. In addition, there is a weak but significant positive correlation between SI_SFG and SI_SFG, suggesting some overlaps in the neurons that encode saliency created by orientation and motion direction.

**Figure 5.**
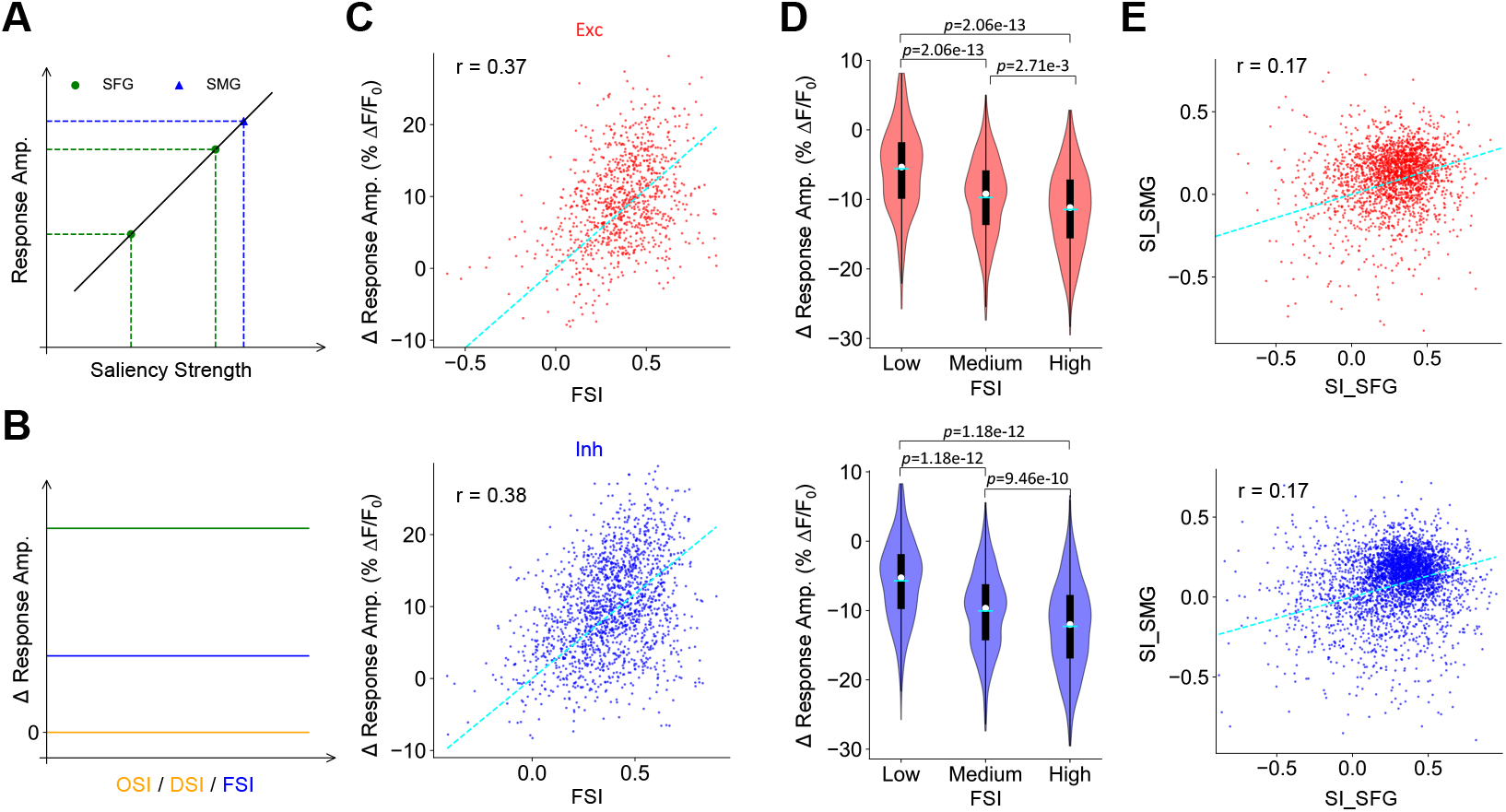
Encoding of visual saliency depends on specific visual features. **A**. For ideal saliency-encoding neurons, response amplitude is a function of saliency strength and is independent of specific visual features. Green dots indicate different orientation contrasts. The SMG could be more or less salient than the SFG. **B**. The response amplitude difference of two salient stimuli should be independent of neurons’ preference for specific orientations, directions, and features. **C**. Difference in response amplitude versus FSI; *r* is Pearson’s correlation coefficient: *p*=3.99e-32 for excitatory neurons, *p*=3.31e-60 for inhibitory neurons. Exc: N=968; Inh: N=1688. Cyan lines are the linear fitting. **D**. The difference in response amplitude elicited by SFG and SMG for saliency-encoding neurons with different levels of feature selectivity. Low: FSI*<*0.25; Medium: 0.25 ≤ FSI*<*0.5; FSI ≥ 0.5. One-way ANOVA, *p*=6.29e-29 for excitatory neurons, *p*=1.27e-83 for inhibitory neurons. Tukey’s range test for pairwise comparisons. **E**. SI_SMG versus SI_SFG; *r* is Pearson’s correlation coefficient: *p*=3.67e-15 for excitatory neurons, *p*=1.80e-24 for inhibitory neurons, Exc: N=2056,Inh: N=3344.

### 2.6 Saliency-encoding neurons are less selective to orientation and motion direction

Neurons tuned to specific visual features, such as orientation and motion direction, form a functional map in the mouse SC [28, 35, 36, 29]. How does this feature encoding relate with saliency encoding? As shown in Fig. 2E and 3D, saliency-encoding neurons are less likely to be selective to orientation or motion direction. To have a more comprehensive understanding of their relationship, for the same group of neurons, we plotted their orientation map, direction map, and saliency maps (Fig. 6A). In the saliency map, the neural coding of saliency strength at different locations is indicated by SI. We found that neurons that show strong orientation or direction selectivity are more likely to have low SI and show weak responses to salient stimuli (Fig. 6A, S3A-C). Indeed, both gOSI and gDSI are negatively correlated with SI for both excitatory and inhibitory neurons (Fig. 6B and C). The weak but significant negative correlation indicates that visual features and saliency are encoded by two groups of neurons that are partially overlapped. This anticorrelation can not be attributed to the response amplitude at the preferred orientation or direction, because feature selectivity is mainly contributed by a decrease at the orthogonal orientation or null direction (Fig.S3D and E).

**Figure 6.**
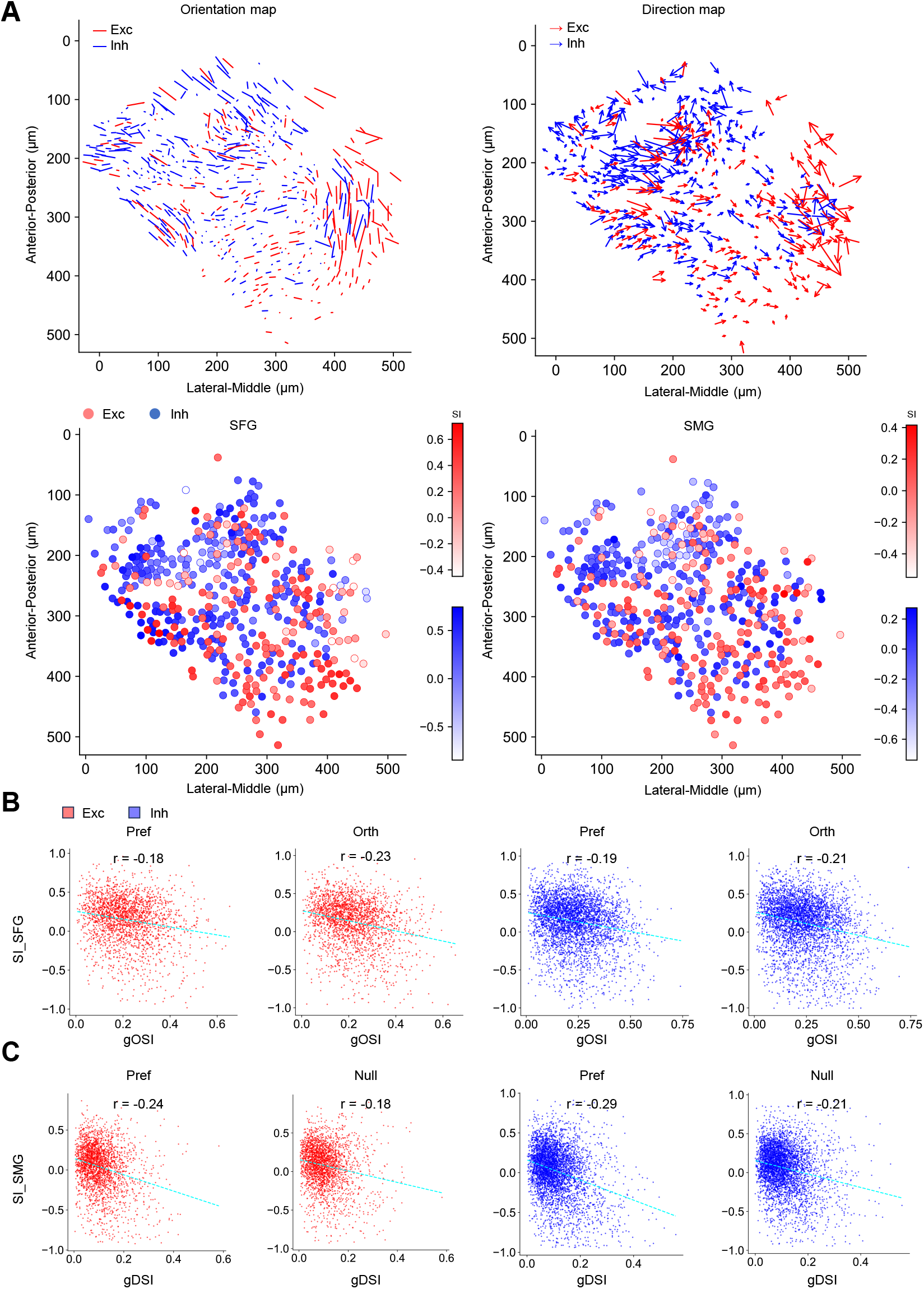
Saliency-encoding neurons are less selective to orientation and motion direction. **A**. Upper panels show the orientation map and direction map represented by excitatory (red) and inhibitory (blue) neurons in an imaging plane. The length of the lines or arrows proportionally correlates with gOSI or gDSI. Lower panels show the saliency map represented by excitatory and inhibitory neurons in the same plane. **B**. SI is anti-correlated with OSI measured by flashing gratings: *r* is Pearson’s correlation coefficient, from left to right: *p*=4.98e-18, 6.34e-28, 7.39e-30, 1.47e-38. Exc: N=1211; Inh: N=2516. **C**. SI is anti-correlated with DSI measured by moving gratings: from left to right: *p*=4.39e-35, 4.23e-20, 9.04e-81, 3.03e-43. Exc: N=2116; Inh: N=3511.

## 3 Discussion

Cracking the neural encoding of visual saliency is important for understanding how useful information is extracted from massive visual input, and such an extracting process is crucial for animals to survive and thrive in an ever-changing environment. In this study, we investigate how excitatory and inhibitory neurons in sSC encode visual saliency made by different visual features.

### 3.1 Main findings

To uncover how visual saliency is encoded in sSC, we applied two-photon calcium imaging to measure the responses of excitatory and inhibitory neurons in awake mice to salient visual stimuli. We found that excitatory and inhibitory neurons show similar response properties in the awake state (Fig. 1). Specifically, a salient stimulus at the RF center of neurons evokes the strongest response amplitude, which decreases as the stimulus moves away from the RF until the distance reaches 40° (Fig. 2 and 3). The response amplitude to salient stimuli is independent of the preferred orientation and motion direction of neurons (Fig. 2 and 3; instead, it is determined by the saliency strength of the stimuli (Fig. 4. However, when the salient stimuli are made by different features, the encoding of visual salience depends on specific visual features (Fig. 5). Thus, the theory proposed by Koch and Ullman [7] is partially supported by experimental evidence in the mouse sSC. Further analysis suggests that saliency-encoding neurons and feature-encoding neurons may belong to different neural ensembles that are partially overlapped (Fig. 6).

### 3.6 Relation to prior work

Prior work indicates that the saliency encoding in V1 depends on neurons’ preference for specific orientation or motion direction [12, 13, 21, 14, 37]. Neurons in cat and monkey V1 are sharply tuned to orientation or motion direction [38, 39]. Their responses to salient stimuli are determined by both the saliency strength of visual stimuli and their preference for orientation or motion direction [12, 13, 21, 14]. In contrast, neurons in the primate sSC are less tuned to these features but show robust responses to salient locations in natural scenes [40, 41, 15], suggesting a feature-agnostic encoding of saliency. In rat sSC, the encoding of visual saliency is not influenced by orientation preference [42]. In the optic tectum, the non-mammalian homologue of SC, neurons encode visual saliency based on motion direction in a preference-independent manner but do not encode orientation-induced saliency [43, 44, 45]. Here, we present direct evidence in mice supporting the notion that saliency encoding in the sSC is independent of both orientation and direction preference but depends on specific visual features. Furthermore, saliency-encoding neurons in mouse sSC are less selective to orientation and motion direction (Fig. 6), echoing similar findings in the primate sSC.

Note that the saliency encoding in the sSC is affected by both anesthesia and cortical inputs [26]. In anesthetized mice, excitatory and inhibitory neurons in the sSC show opposite responses to motion contrast, with both types being suppressed by the orientation contrast [25]. However, in the awake state, our study demonstrates that excitatory and inhibitory neurons show similar responses to salient stimuli, consistent with recent findings showing strong responses to the edge of drifting or flashing gratings in both types of neurons [46].

Moreover, the saliency encoding is also influenced by the target size. In our study, the maximal response of sSC neurons is elicited by a 10° salient stimulus at the RF center, indicating their role as saliency detectors. However, when the target size exceeds the RF size, iso-feature suppression at the center causes the maximal response to be elicited by the edge, leading to neurons behaving as edge detectors [46].

### 3.3 Neural implementation of visual saliency

In primate V1, encoding of visual saliency depends on the neuron’s preference for specific orientations or motion directions. For a cortical neuron, when the orientation surrounding its RF matches its preferred orientation, iso-orientation stimuli within its RF elicit stronger responses than cross-orientation stimuli. Conversely, when the surrounding orientation is orthogonal to its preferred orientation, cross-orientation stimuli are more effective [12]. One can understand such contextual effects by convolving the visual stimulus with its subthreshold RF (Fig. **??**), which reflects the integrated synaptic inputs and is larger than the spike RF [47].

However, the convolution fails to predict the preference-independent encoding of visual saliency within features in the mouse sSC, which might be addressed by integrating long-range inhibition as proposed by the model. This long-range inhibition could be contributed by local inhibitory neurons with horizontally extended dendritic arbors reaching up to 500 *µ*m, accounting for ∼60% of all neurons in the sSC [34, 31]. In contrast, inhibitory neurons in V1 exhibit heterogeneous morphologies and represent only ∼25% of the total neuronal population [48]. Another contributing factor could be functional patches in the sSC [28, 29]: first, the size of these patches is comparable to the observed long-range effect over ∼40° (Fig. **??**A and **??**C); second, the connection strength between neurons is affected by both their distance and functional similarity [49, 50, 51]. Additionally, wide-field excitatory neurons with large dendrites may also play a role in saliency encoding [34]. Future modeling work may examine how these factors collectively contribute to the feature-independent saliency map in the sSC.

The neural encoding of visual saliency in the sSC could be contributed by direct inputs from the retina and V1. In the retina, salient stimuli made by orientation or motion direction elicit stronger responses in retinal ganglion cells (RGCs) compared to uniform stimuli [52, 53]. Importantly, the robust response of object motion sensitive (OMS) RGCs to salient moving stimuli is independent of direction preference [54, 55], indicating that saliency encoding in sSC could be partially inherited from the retina, if not entirely. In contrast, cortical inputs from V1 reduce the distinction between responses evoked by cross-orientation and iso-orientation stimuli, thereby compromising the encoding capacity [26]. This implies that saliency encoding in the sSC is less likely to rely on inputs from V1.

### 3.4 The role of superior colliculus in visual attention

In primates, the sSC plays a critical role in bottom-up attention and is not involved in arousal-related top-down attention [15, 19]. Supporting this notion, our data demonstrated that saliency encoding is not affected by pupil size (Fig. S1E), a common indicator of an animal’s arousal state [56]. The sSC projects to many brain regions, including the pulvinar, the lateral geniculate nucleus, and the intermediate layer of SC (iSC) [34, 57]. Additionally, iSC also receives inputs from the prefrontal cortex carrying top-down information [58], and neurons in iSC respond to stimuli from different sensory modalities [59]. Thus, iSC may serve as an ideal hub to integrate the feature-independent saliency map from sSC with goal-driven attention to generate a modality-independent priority map that determines the attended location. Indeed, inactivating deep layers of the primate SC compromises the behavioral performance in visual-attention tasks [60]. One exciting research direction is to explore how the feature-independent visual saliency map in sSC is transformed into the modality-independent priority map in iSC.

## 4 Materials and Methods

All experimental procedures were performed under the animal welfare guidelines and approved by the Institutional Animal Care and Use Committee at the Chinese Institute for Brain Research, Beijing.

### 4.1 Animal

Vglut2-ires-Cre (JAX no. 028863) or Vgat-ires-Cre (JAX no. 028862) mice were crossed with Ai14 mice (JAX no. 007914) to express tdTomato in either excitatory (Vglut+) or inhibitory (Vgat+) neurons. One male and two female Vglut2-tdTomota mice and four female Vgat-tdTomato mice were used at ages 2-4 months.

### 4.2 Viral injection

We injected adeno-associated virus (AAV) expressing non-floxed GCaMP8m (AAV2/9-syn-jGCaMP8m-WPR) into the SC of Vglut2-tdTomato and Vgat-tdTomato mice. After 2–3 weeks, we surgically implanted a cranial window coupled to a transparent silicone plug to expose the posterior-medial portion of the sSC, corresponding to the upper-temporal part of the visual field. After 3 days, we used two-photon microscopy to image calcium signals in the sSC of head-fixed awake mice.

### 4.3 *In vivo* two-photon calcium imaging

The animal was head-fixed on a treadmill and free to move. Following a 15-minute habituation period, two-photon imaging was performed using a microscope (Ultima 2P Plus, Bruker) with a 16×, 0.8 NA, 3 mm WD objective (CF175, Nikon). A tunable femtosecond laser (InSight X3+ Dual, Spectra-Physics) was raster scanned by resonant galvanometers. GCaMP8m and tdTomato were excited at 920 nm, with the laser power at the sample plane typically set between 30−50 mW. A 550 *µ*m ×550 *µ*m field of view was scanned at 10 Hz as a series of 512 pixels × 512 pixels images, and the imaging depth was up to 350 *µ*m. Emitted green and red light were split with a dichroic mirror (t565lpxr), passed through two bandpass filters (et525/70m-2p and et595/50m-2p), and detected by two GaAsP tubes (H10770PB-40, Hamamatsu). The animal’s locomotion, pupil size, and positions were recorded and synchronized with the image acquisition. During the imaging session, the mice exhibited only rare eye movements and locomotion.

### 4.4 Visual stimulation

A 21-inch LED monitor was placed 17 cm away from the mouse’s right eye, centered at 95° azimuth and 25° elevation, covering a visual field of 104°× 80°. To investigate how visual saliency is encoded in the sSC, we presented six types of visual stimuli in a 40°× 60° space that effectively covered the visual field recorded during each imaging session. **(1)** 4×6 (one at a time) 10° × 10° flashing squares (1 s black or white + 1 s grey) to map the receptive field (RF). **(2)** A full-field flashing square grating with a spatial frequency of 0.1 cycles per degree in 6 orientations and 4 phases (4 s grating + 1 s grey) to measure the orientation selectivity. **(3)** 4×6 (one at a time) 10° ×10° flashing square gratings in vertical (0°) or horizontal (180°) orientations on a grating background in the orthogonal orientation. **(4)** A 10°×10° flashing square grating in 6 orientations at the RF center with a vertical or horizontal grating background. **(5)** A full-field moving square grating with a spatial frequency of 0.1 cycles per degree and a temporal frequency of 2 Hz in 12 directions measures the direction selectivity (2 s grating + 1 s grey). **(6)** 4×6 (one at a time) 10°×10° moving gratings in 0°, 90°, 180°, and 270° directions on a grating background moving in the opposite directions. All stimuli were displayed for 10 repetitions, and the sequence was pseudo-randomized for each stimulus.

### 4.5 Data analysis

#### 4.5.1 Measurement of calcium responses

Brain motion during imaging was corrected using NoRMCorre [61]. Regions of interest (ROIs) were manually drawn using the Cell Magic Wand Tool (ImageJ) and fitted with an ellipse in Python. Fluorescence traces of each ROI were extracted after estimating and removing contamination from surrounding neuropil signals as described previously [29, 62, 63]. The true fluorescence signal of a neuron was calculated as *F*_true_ = *F*_raw_ − (*r* ·*F*_neuropil_), where *r* is the out-of-focus neuropil contamination factor, estimated to be ∼0.7 for our setup. Slow baseline fluctuations were removed using detrended fluctuation analysis with a 15-second sliding window.

For any given stimulus, the response of a neuron was defined by the fluorescence trace in its ROI during the stimulus period:

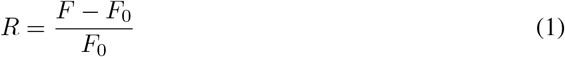

where *F* is the instantaneous fluorescence intensity and *F*_0_ is the mean fluorescence intensity without visual stimulation (grey screen).

Two criteria were applied to interpret ROIs as neurons: 1) The ROI size was limited to 10-20 *µ*m to match the size of a neuron; 2) The ROI response had to pass a signal-to-noise ratio (SNR) of 0.5 [33]:

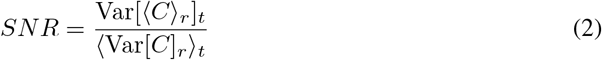

where *C* is the *N*_*t*_ (time samples) × *N*_*r*_ (stimulus repetitions) response matrix, *t* = 1, …, *N*_*t*_ and *r* = 1, …, *N*_*r*_, ⟨·⟩_*r*_ and ⟨·⟩_*t*_ are the means over repetitions or time respectively, and Var[·]_*r*_ and Var[·]_*t*_ are the corresponding variances. All ROIs meeting these criteria were selected for further analysis, yielding a total of 7020 neurons, including 2565 excitatory neurons and 4455 inhibitory neurons.

### 4.5.2 Quantification of neuronal responses

To quantify the orientation tuning, we calculated the orientation selectivity index (OSI) as the normalized difference between the amplitude at the preferred and orthogonal orientations and the global orientation selectivity index (gOSI) as the normalized amplitude of the response-weighted vector sum of all orientations:

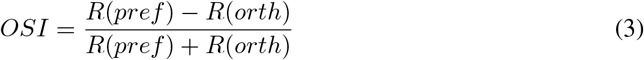

Where *R*(*pref*) *R*(*orth*) are the average response to the preferred and orthogonal orientation, respectively.

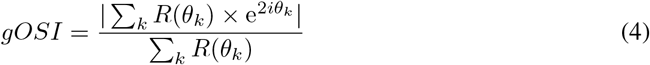

Where *θ*_*k*_ is the *k*^th^ orientation in radians and *R*(*θ*_*k*_) is the average response during the stimulus period at that orientation.

To quantify the tuning of a neuron to motion directions, we calculated the direction selectivity index (DSI) as the normalized amplitude of the response-weighted vector sum of all directions:

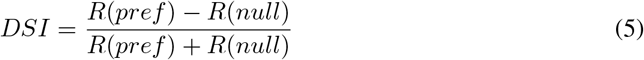

Where *R*(*pref*) *R*(*null*) are the average response to the preferred and null motion direction, respectively.

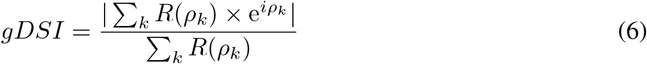

Where *ρ*_*k*_ is the *k*^th^ direction in radians and *R*(*ρ*_*k*_) is the average response during the stimulus period at that direction.

To quantify the preference for salient stimuli, we calculated the saliency index (SI) as the relative difference between its response to SFG or SMG and that to the background.

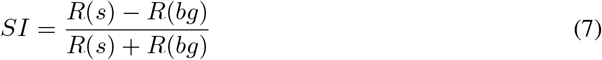

Where *R*(*s*) is the average response to a salient stimulus during the stimulus period, and *R*(*bg*) is the average response to the background.

To quantify the preference for specific visual features, we calculated the feature selectivity index (FSI) as the relative difference between its response to flashing gratings and moving gratings.

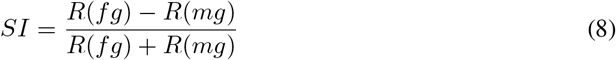

Where *R*(*fg*) is the average response to flashing gratings at the preferred orientation, and *R*(*mg*) is the average response to moving gratings at the preferred motion direction.

To quantify RF size and locate its center, calcium responses at 4×6 locations were fitted with a 2-D Gaussian function (Equation 9),

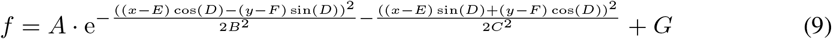

The RF size is defined as the area at half maximum, which equals *π* · 2 ln 2 · *B* · *C*. We omitted analysis of the RF if the coefficient of determination for this fit was below 0.5. We also calculated the RF center as the center of mass for evoked responses and found consistent results with both approaches.

### 4.6 Neural modeling

For simplicity, the model was built in 1-D visual space with only two stimuli: positive and negative. Functional patches are simulated with a sinusoidal function, termed the functional preference index (FPI), which quantifies the preference of each neuron at different spatial locations. Since the OSI is mainly correlated with the response at the orthogonal orientation (Fig. S3D), we set a neuron’s response amplitude to its preferred stimulus as 1, while its responses to non-preferred stimuli varied with FPI.

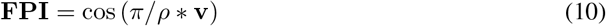

Where *ρ* denotes the patch size set to 30°, and **v** is a vector representing the visual space of [-60°, 60°].

The RF of each neuron was simulated with a 1-D Gaussian function.

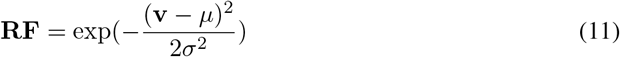

Where *µ* is the location of the RF center in visual space, and *σ* is set to 5° for the subthreshold RF. A neuron’s response to a given visual stimulus is simulated by convolving its RF with the stimulus.

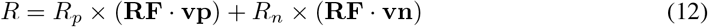

Where **vp** and **vn** are the positive and negative stimuli, and *R*_*p*_ and *R*_*n*_ are the response amplitude to the positive and negative stimuli.

### 4.7 Statistics

The Kolmogorov–Smirnov test was used to assess the normality of data distributions. Parametric tests were applied to normally distributed data, while non-parametric tests were used for all other data. Statistical significance was defined as p<0.05. Sample sizes were not predetermined by statistical methods but were based on common practices in the field.

## 5 Supplement

Here we report details related to the Results, Discussion, and Methods sections.

**Figure S1:**
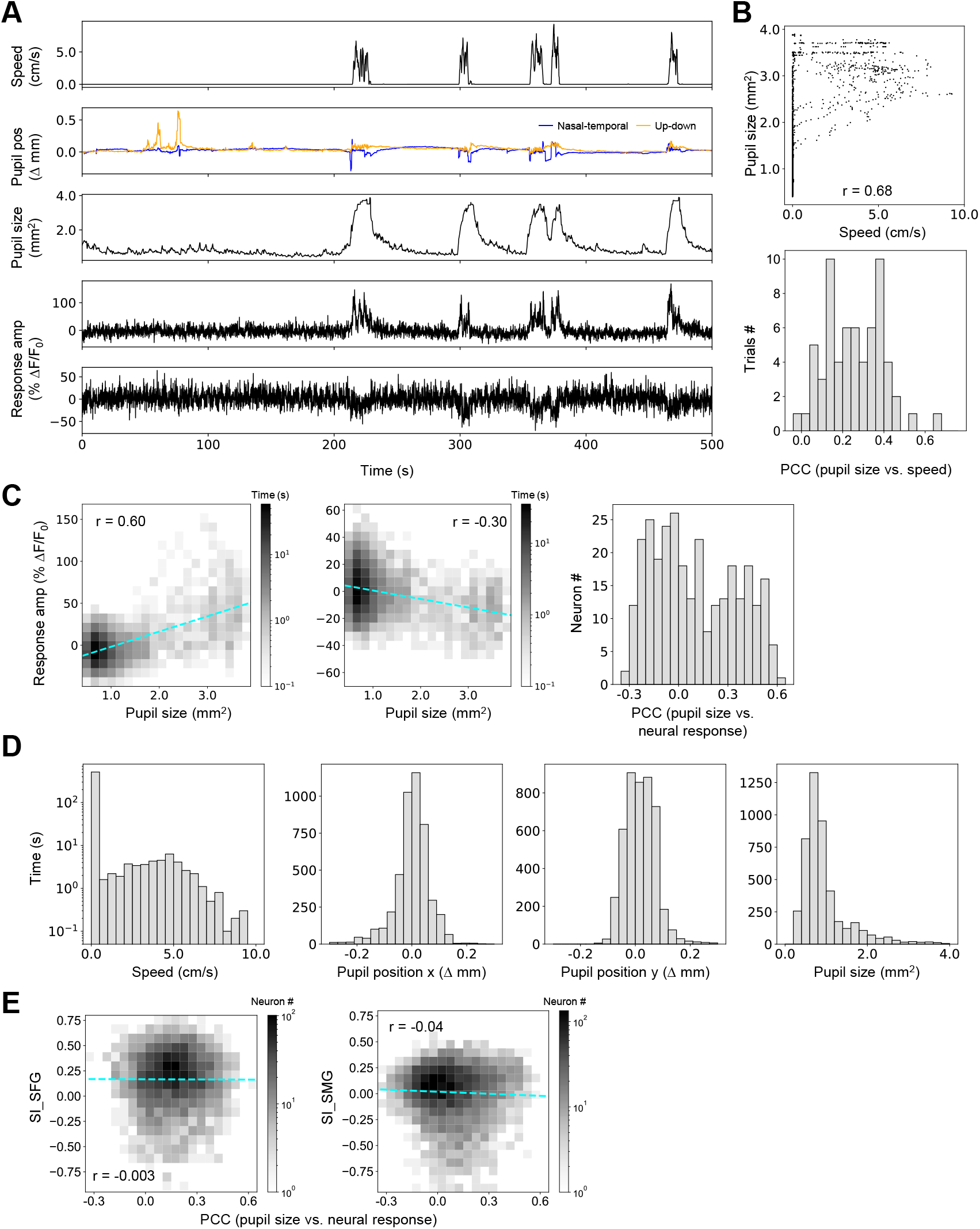
Behavioral modulation of neuronal responses to salient stimuli (related to Fig. 1). **A**. Locomotion speed, pupil positions, pupil size, and calcium responses in an example mouse. **B**. Top, Pupil size versus locomotion speed: *r* denotes the Pearson’s correlation coefficient (PCC). Bottom, histogram of their PCC. **C**. Left, calcium responses of two example neurons versus pupil size. Cyan lines indicate the linear fitting. Right, histogram of their PCC. **D**. Histograms of locomotion speed, pupil positions, and pupil size for all animals. **E**. SI_SFG and SI_SMG versus PCC between pupil area and neural responses.

**Figure S2:**
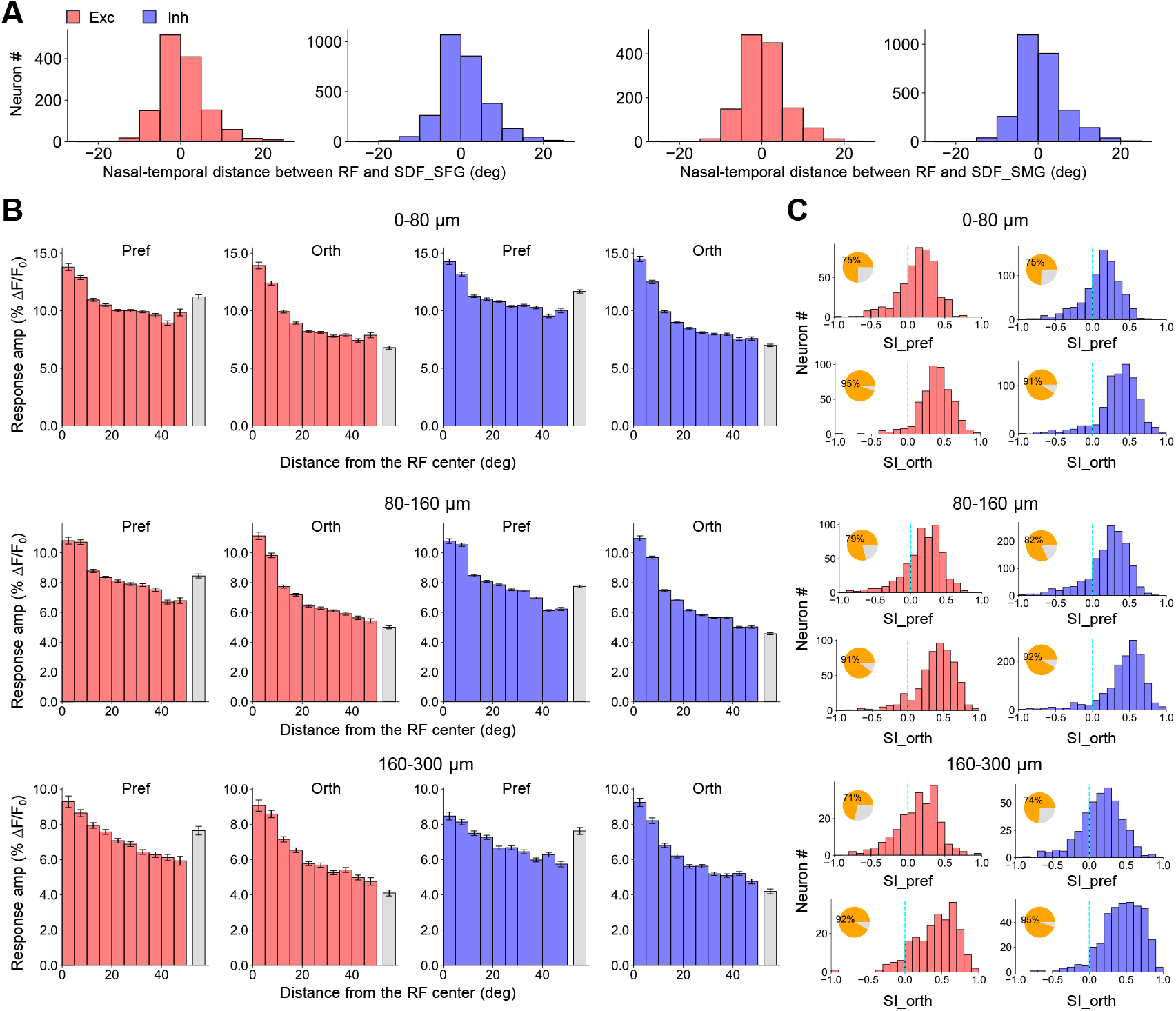
Preference- and depth-independent saliency encoding in the mouse sSC (related to Fig. 2). **A**. The relative distance along the nasal-temporal axis between the RF center and the center of the saliency detection field (SDF). The center of SDF is defined as the center of mass for SFG- or SMG-evoked responses (SDF_SFG or SDF_SMG). **B**. Response amplitude to SFG and the background grouped by orientation preference across the depth. Exc: N=498, 563, 212; Inh: N=817, 1464, 403. **C**. Histograms of SI across the depth.

**Figure S3:**
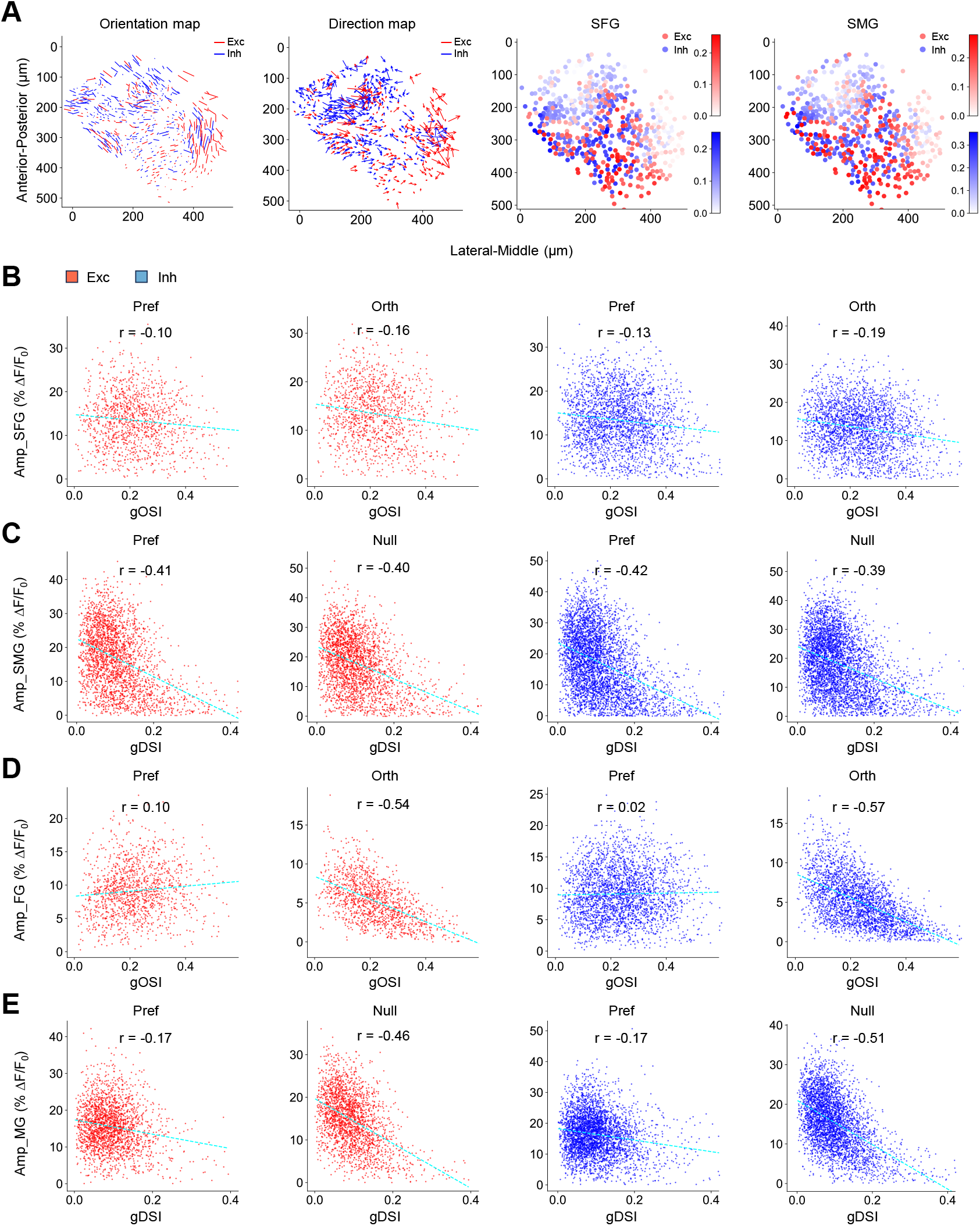
Neurons tuned to the orientation and motion direction show weaker responses to salient stimuli compared to non-tuned neurons. **A**. Left panels show the orientation map and direction map represented by excitatory (red) and inhibitory (blue) neurons in an imaging plane (the same as Fig.**??**A). The right part shows the response amplitude to SFG and SMG of the same group of neurons. **A**. Response amplitude to SFG is anti-correlated with OSI: r is Pearson’s correlation coefficient, *p*<0.001 for all correlations. **C**. Response amplitude to SMG is anti-correlated with DSI: *p*<0.001 for all correlations. **D**. Response amplitude to flashed gratings (FG) at the preferred and orthogonal orientations versus OSI: *p*<0.01, <0.001, <0.05, <0.001 for correlations from left to right. **E**. Response amplitude to moving gratings (MG) at the preferred and null directions versus DSI: *p*<0.001 for all correlations.

**Figure S4:**
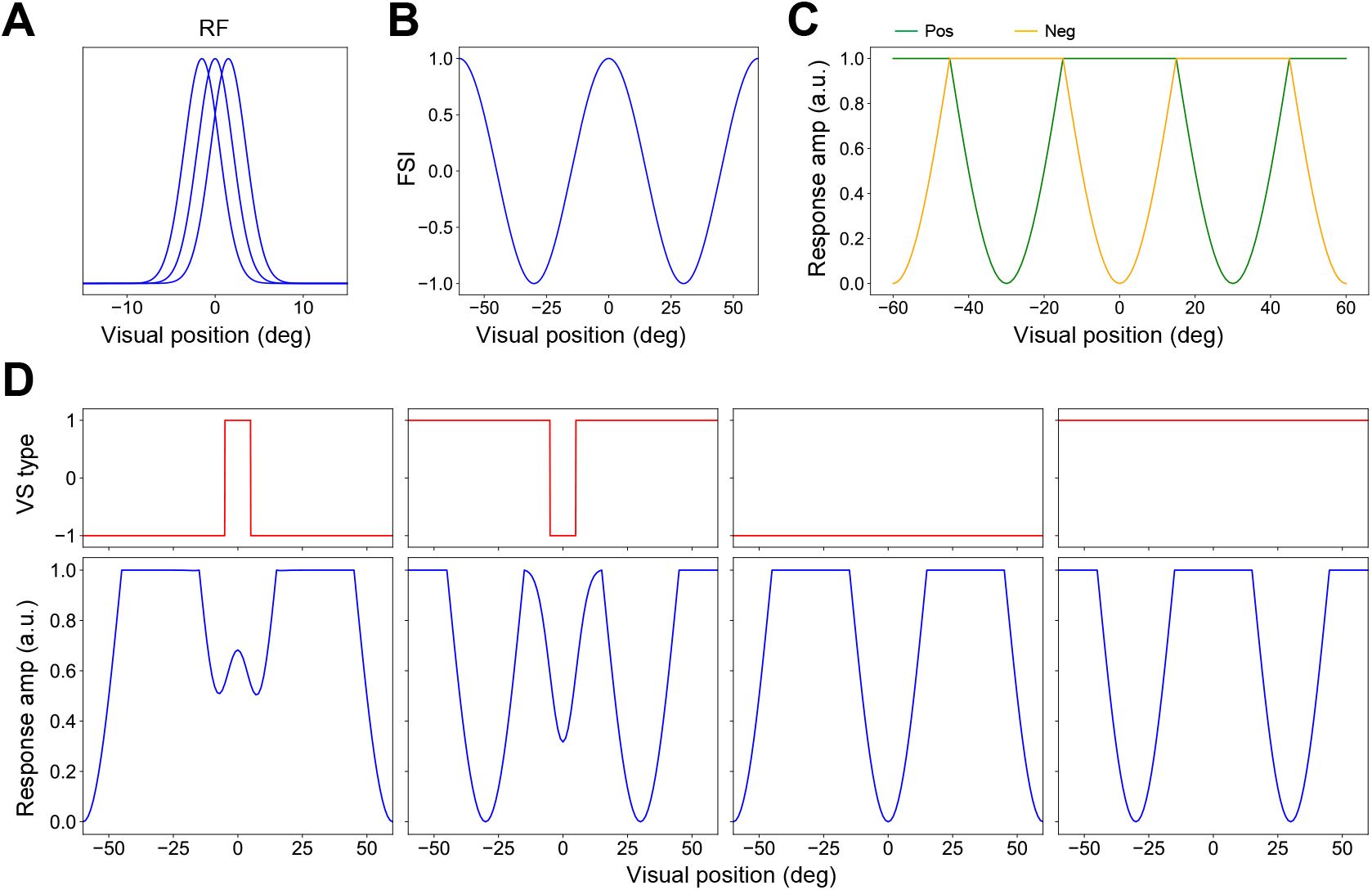
Neural modeling for feature-dependent saliency in V1. **A**. Synaptic RFs for three nearby neurons. **B**. FPI of neurons in 1-D visual space. **C**. Neuronal responses to positive and negative stimuli. **D**. Top, four visual stimuli. Bottom, simulated neuronal responses in the visual space.

## Acknowledgments

Y.-t.L. is supported by startup funds from CIBR, the National Natural Science Foundation of China (32271060), and the Beijing Natural Science Foundation (IS23073). L.-y.L. is supported by the Beijing Natural Science Foundation (5244028).

## Author contributions

Y.-t.L. supervised the project; Y.-t.L. and R.W. designed the experiments; R.W. collected all the data; Y.-t.L., R.W., J.X., and C.L. analyzed the data; Z.Z. and Y.-t.L. carried out the neural modeling; R.W. and Y.-t.L. prepared figures; Y.-t.L. and L.-y.L. wrote the manuscript.

## Data availability

Data will be available in a public repository upon acceptance of the manuscript.

## Code availability

Code will be available in a public repository upon acceptance of the manuscript.

